# *Enterococcus faecalis* is involved in the progression of the early stages of latent chronic pancreatitis surrounding pancreatic cancer tissue

**DOI:** 10.64898/2026.07.20.739687

**Authors:** Kaho Nishikori, Shinji Takamatsu, Munefumi Shimosaka, Risa Uemura, Yudai Ishida, Rikako Sugawa, Muya Matsumoto, Asuka Ogata, Daisuke Sakon, Motoharu Inui, Daisaku Yamada, Hirofumi Akita, Jumpei Kondo, Takahiro Kodama, Yoshihiro Kamada, Hidetoshi Eguchi, Eiichi Morii, Eiji Miyoshi

## Abstract

(Objective) In our previous research, we identified latent chronic pancreatitis in the normal tissue surrounding pancreatic cancer. We also discovered the presence of *Enterococcus faecalis* (*E. faecalis*), a type of intestinal bacterium, in the pancreatic fluid and tissue of pancreatic cancer patients, suggesting it may be one of the factors contributing to the development of latent chronic pancreatitis. In this study, we performed pathological analyses to investigate its characteristics and investigate a possibility of *E. faecalis* infection. (Methods) Pathological analyses were performed, using 16 cases of pancreatic cancer and intraductal papillary mucinous neoplasia (IPMN) involving lesions in the pancreas tail. The involvement of *E. faecalis* was investigated with immunohistochemical analysis and serological methods. (Results) All cases exhibited inflammatory changes in pancreatic tissue without a clinical diagnosis of chronic pancreatitis, along with macrophage infiltration. These changes did not significantly differ according to preoperative treatment. DNA encoding *E. faecalis* 16s ribosomal RNA was detected in many cases, however, a positive immunostaining to *E. faecalis* was observed in only a few cases. Serum capsular polysaccharide (CPS) antibody levels exceeding the mean values were observed in patients with established chronic pancreatitis, while the level was not correlated with *E. faecalis* immunostaining. (Conclusion) These results suggest the *E. faecalis* infection is involved in the early stage of the progression of chronic latent pancreatitis and the diagnostic technology incorporating novel multi-biomarkers may be useful for identifying high-risk individuals for future pancreatic cancer development.

## Introduction

Pancreatic ductal adenocarcinoma (PDAC) is a highly aggressive malignancy with a 5-year survival rate below 10%, and many cases are already locally advanced or metastatic at diagnosis ^1,2^. For true early detection of pancreatic cancer, it is essential to define the initial stages of carcinogenesis, i.e., “precancerous lesions,” and to establish indicators for early intervention. Previous pathological studies have identified pancreatic intraepithelial neoplasia (PanIN) as a primary precursor lesion in pancreatic cancer development ^3^. Among PanIN classifications, PanIN-3 has been considered the lesion immediately preceding invasive cancer. However, recent studies have reported the intriguing finding that only a small fraction of PanIN lesions acquire additional alterations and progress to invasive cancer ^4^.

In our previous study, we performed pathological analysis of peritumoral tissues from autopsy specimens of pancreatic cancer and found that pancreatic tissues showing histological features of chronic pancreatitis were frequently observed, even in patients without clinical symptoms ^5^. This finding suggests that chronic pancreatitis may serve as a high-risk environment for pancreatic cancer development. In this study, we refer to this condition as “latent chronic pancreatitis.” In chronic pancreatitis, persistent inflammation and fibrosis may induce microductal changes that promote the formation of PanIN and intraductal papillary mucinous neoplasia (IPMN). Possible causes of latent chronic pancreatitis include age-related pancreatic steatosis, bacterial infections, and immune responses triggered by bacterial components. Indeed, fatty pancreas has increasingly been recognized as a risk factor for pancreatic cancer ^6^.

Meanwhile, an increasing number of studies have reported associations between pancreatic cancer and bacterial infection ^7–9^. We have long focused on the potential role of bacterial infection in pancreatic diseases. Using postoperative pancreatic drainage fluid and pancreatic tissue from patients with pancreatic cancer, we detected abundant bacterial 16S rRNA genes in pancreatic fluid. Species-level PCR and immunostaining for Enterococcus spp. revealed that *Enterococcus faecalis* (*E. faecalis*) was frequently detected ^10^. Furthermore, we have recently reported that highly pathogenic *E. faecalis* may contribute to the pathogenesis of pancreatitis by invading pancreatic ductal epithelial cells ^11^. Oral administration of this strain in mice induced pathological changes resembling chronic pancreatitis ^12^.

However, in our previous pathological analysis ^5^, we could not exclude the possibility of postmortem pancreatic degeneration or obstructive pancreatitis caused by pancreatic cancer. Therefore, in the present study, we focused on cancers and IPMN located in the pancreatic tail and examined whether latent chronic pancreatitis is present in the pancreatic head region immediately proximal to the lesion, where obstructive effects are unlikely. In addition, pathological and serological analyses were performed to investigate the involvement of *E. faecalis* in these patients.

## Materials and methods

### Clinical specimens

This retrospective study was conducted in accordance with the Declaration of Helsinki and was approved by the Institutional Review Board of the University of Osaka Hospital (reference numbers 14107 and 15212). De-identified specimens and data were accessed for research purposes between 01/04/2024 to 31/03/2026. Residual specimens of pancreatic cancer and adjacent normal pancreatic tissue resected at the Department of Gastroenterological Surgery in the University of Osaka Hospital, were used. The targeted diseases were pancreatic tail cancer and IPMN of the pancreatic tail. Normal pancreatic tissue at least 3 cm proximal to the tumor was used for pathological analysis. S1 Table summarizes the clinical information of all pancreatic tissue samples. Corresponding patient serum samples were stored at-80°C until use. In addition to the 16 patient serum samples described above, serum samples from 169 patients with chronic pancreatitis (Ogaki Municipal Hospital and JCHO Osaka Hospital), 203 patients with pancreatic cancer (the University of Osaka Hospital), 91 healthy volunteers of medical check-up people whose liver/pancreatic functions were within normal ranges (aMs new Ohtani Clinic) were analyzed.

### Histological and Immunohistochemical Staining

Formalin-fixed, paraffin-embedded pancreatic tissue sections (4 μm) were stained with hematoxylin and eosin (H&E) using routine procedures and examined by light microscopy.

For immunohistochemical staining, deparaffinized and rehydrated sections were subjected to heat-induced antigen retrieval in sodium citrate buffer (pH 6.0) at 95°C, followed by blocking of endogenous peroxidase with 1% hydrogen peroxide and nonspecific binding with 5% normal donkey serum in TBST. Sections were incubated with primary antibodies overnight at 4°C and with horseradish peroxidase–conjugated secondary antibodies for 30 minutes at room temperature. Signals were developed using a DAB detection kit (Nacalai Tesque, Kyoto, Japan), and nuclei were counterstained with hematoxylin. Antibodies used are listed in S2 Table.

For double immunofluorescence staining, sections were processed for antigen retrieval as described above and blocked with 5% normal donkey serum in TBST. Next, rabbit anti-human CD163 antibody was reacted, washed, and incubated with CF488-labeled donkey anti-rabbit secondary antibody (Biotium, Fremont, CA). After further washing, the samples were incubated with DAPI and fluorescently labeled rabbit anti-human CD68 antibody (HiLyte Fluor 555, DOJINDO, Kumamoto, Japan). The slides were mounted with aqueous mounting medium (Aqua-Poly/Mount, Polysciences, Warrington, PA) and images were acquired using an HS All-in-One Fluorescence Microscope BZ-X810 (KEYENCE, Osaka, Japan).

### Blinded histological assessment of H&E-stained slides

For each of the 16 cases, 10 fields were randomly selected and photographed at 400 × magnification from H&E-stained sections using an HS All-in-One Fluorescence Microscope BZ-X810 (KEYENCE). Cases were anonymized and randomly numbered. Image files and evaluation sheets were distributed to five expert pancreatic pathologists, who independently assessed each image according to predefined criteria.

The following features were evaluated: (1) fatty change of the pancreas, (2) fibrosis, and (3) immune cell infiltration. Each feature was scored on a 4-point scale: 0 (none, 0–5%), 1 (mild, 6–33%), 2 (moderate, 34–66%), and 3 (severe, 67–100%), yielding a maximum of 30 points per case for each item. Mean scores were calculated to obtain the fatty pancreas score, fibrosis score, and immune cell infiltration score for each case. To standardize the evaluation, representative reference images were provided for each category. During the first round of assessment, re-examination and correction of scores were not permitted. After all 160 images had been scored, evaluators were allowed a second round to review the images and revise their scores.

### Quantification of Immunohistochemical Positivity

DAB-stained images for each antibody were analyzed using Fiji (Fiji is just ImageJ) ^13^. Color deconvolution and image binarization were performed according to previously published methods ^14,15^. The hematoxylin-positive and DAB-positive areas were measured separately, and the ratio of DAB-positive area to hematoxylin-positive area was defined as the immunostaining positivity score. For comparative analyses, upper and lower threshold values were standardized across all images included in the comparison. Mean CD68 and CD3 positivity was calculated as the average positivity score across 10 randomly selected fields per case at 100× magnification.

### DNA extraction and PCR amplification of 16S rRNA gene for bacterial and *E. faecalis* detection

Genomic DNA was extracted from pancreatic tissue using the DNeasy® Blood & Tissue Kit (QIAGEN, Hilden, Germany) according to the manufacturer’s instructions. The extracted DNA was used for PCR amplification of the 16SrRNA gene for bacterial detection and E. faecalis identification using T100 thermal cycler (BioRad, Hercules, CA). GoTaq Green Master Mix (Promega, Madison, WI) was used for PCR amplification of the target gene. The primers for the bacterial 16S rRNA gene (V3-V4 region) were as follows; SD-bact-0341b-S-17: 5’-CCTACGGGNGGCWGCAG-3’ SD-bact-0785-a-A-21: 5’-GACTACHVGGGTATCTAATCC-3’ After an initial denaturation at 94°C for 5 min, the PCR reaction consisted of three steps: 30 cycles of denaturation at 94°C for 40 s, annealing at 55°C for 2 min, and extension at 72°C for 1 min, followed by a final extension at 72°C for 7 min.

The primers for the DNA encoding *E. faecalis* 16S rRNA were Faecalis2-Ent151F: 5’-ACACTTGGAAACAGGTGC-3’, and Faecalis2-Faecal449R: 5’-AGTTACTAACGTCCTTGTTC-3’.

After an initial denaturation at 94°C for 3 min, the PCR reaction consisted of three steps: 40 cycles of denaturation at 95°C for 30 s, annealing at 55°C for 30 sec, and extension at 72°C for 25 sec, followed by a final extension at 72°C for 7 min. The amplified products were electrophoresed on a 2% agarose gel (in 1x TAE buffer), stained with Midori Green Direct (NIPPON-Genetics, Tokyo, Japan), and visualized with a transilluminator.

### Human CPS-ELISA (Enzyme-Linked ImmunoSorbent Assay)

Preparation of CPS from E. faecalis and ELISA of CPS antibody in serum were performed according to our previous methods ^10^ with slight modification.

### Statistical analysis

Statistical analysis was performed using Microsoft Excel® 2013 (Microsoft, Redmond, WA) JMP Student Edition 18 (SAS Institute, Cary, NC) Data are expressed as mean ± SD, and p < 0.05 was considered statistically significant.

## Results

### Latent chronic pancreatitis in histologically normal pancreatic tissue proximal to pancreatic tail cancer

Pathological analysis was performed on 16 cases without a clinical diagnosis of chronic pancreatitis. Histologically normal pancreatic tissue located at least 3 cm proximal to pancreatic tail tumors, including pancreatic ductal adenocarcinoma and IPMN, as examined in each case. Representative H&E staining is shown in Fig 1A. As expected, fatty degeneration, fibrosis, and inflammatory cell infiltration, which are characteristic pathological features of chronic pancreatitis, were observed in all cases. Blinded evaluation by multiple pathologists showed no differences in fatty degeneration, fibrosis, or inflammatory cell infiltration according to the type of preoperative treatment for pancreatic cancer (Fig 1B). Correlation analysis between these pathological scores and clinical parameters, including pancreatic enzyme levels and HbA1c, revealed significant correlations between BMI and fatty degeneration score, and between fibrosis score and immune cell infiltration score (Fig 1C).

**Fig 1.**
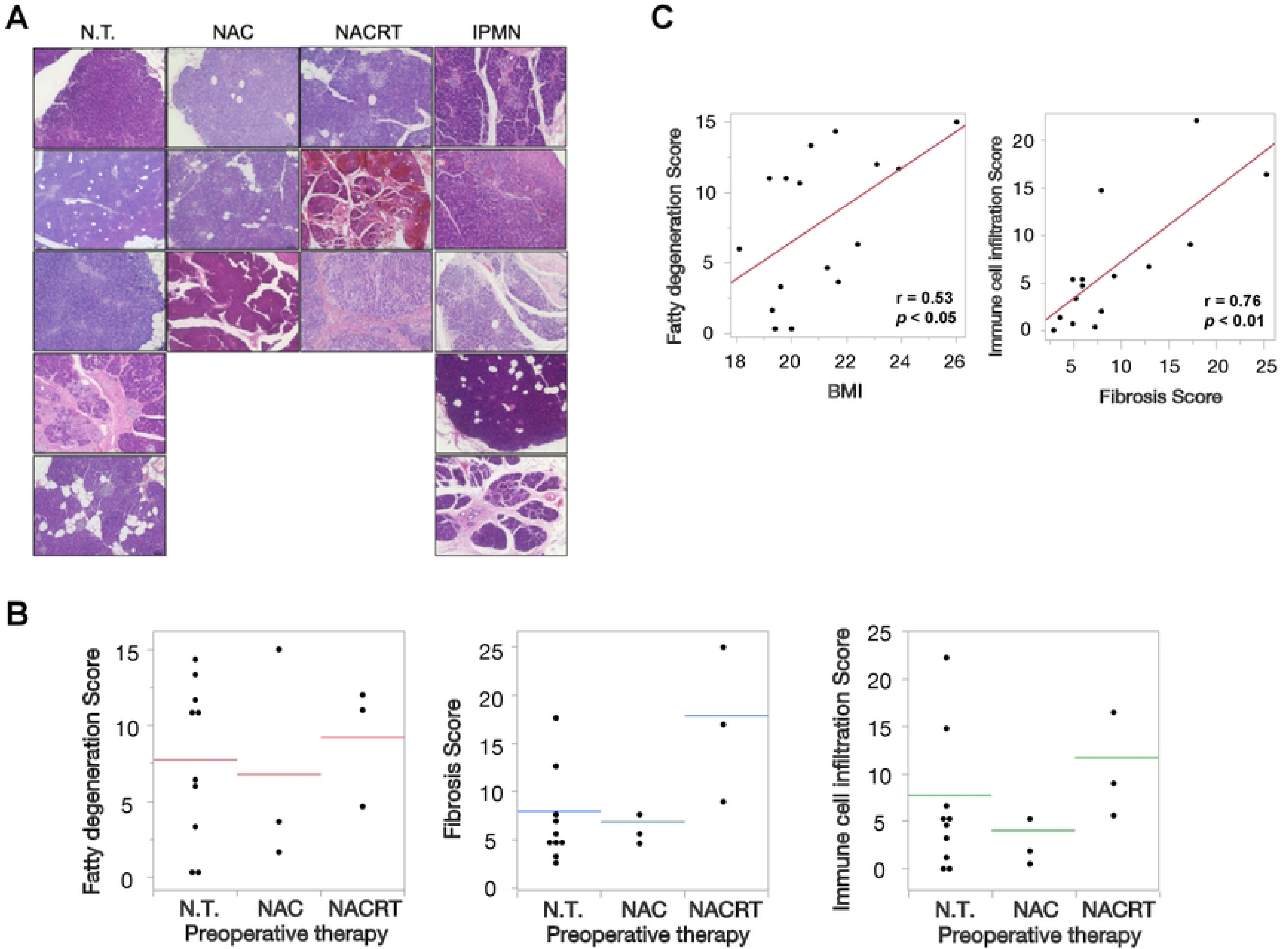
Latent chronic pancreatitis in histologically normal pancreatic tissue proximal to pancreatic tail cancer A. Representative H&E staining of pancreatic tissue from 16 cases of pancreatic tail cancer and pancreatic tail IPMN (40× magnification). Scale bars = 50 µm N.T., Pancreatic tail cancer without preoperative therapy; NAC, Pancreatic tail cancer with preoperative chemotherapy; NACRT, Pancreatic tail cancer with preoperative chemoradiotherapy; IPMN, Intraductal papillary mucinous neoplasm. B. Blinded pathological scores for fatty degeneration, fibrosis, and immune cell infiltration according to preoperative therapy (bars indicate mean values). C. Left: Pearson correlation between blinded fatty pancreas score and patient BMI. Right: Pearson correlation between blinded fibrosis score and immune cell infiltration score.

### Macrophage-dominant inflammatory infiltrates in latent chronic pancreatitis

To characterize the immune cells observed in Fig 1A, immunohistochemical staining of pancreatic tissue was performed using various antibodies. High-magnification images are shown in Fig 2A, and low-magnification images are provided in S1A Fig. Based on the significant correlation between fibrosis and immune cell infiltration observed in Fig 1C, two regions were selected for comparison: the parenchyma-fibrosis interface with prominent immune cell infiltration (Fig 2A left) and areas of dense fibrosis (Fig 2A right). In both regions, CD68-positive macrophages were the most abundant cells among the markers examined (S1B Fig). CD3-positive T cells did not infiltrate the densely fibrotic areas but accumulated focally within the parenchyma (Fig. 2A, upper panels). After exclusion of nonspecific staining, αSMA-positive activated pancreatic stellate cells were mainly located within fibrotic areas, with more prominent activation at the parenchyma-fibrosis interface than in areas of severe fibrosis (Fig 2A, lower panels).

**Fig 2.**
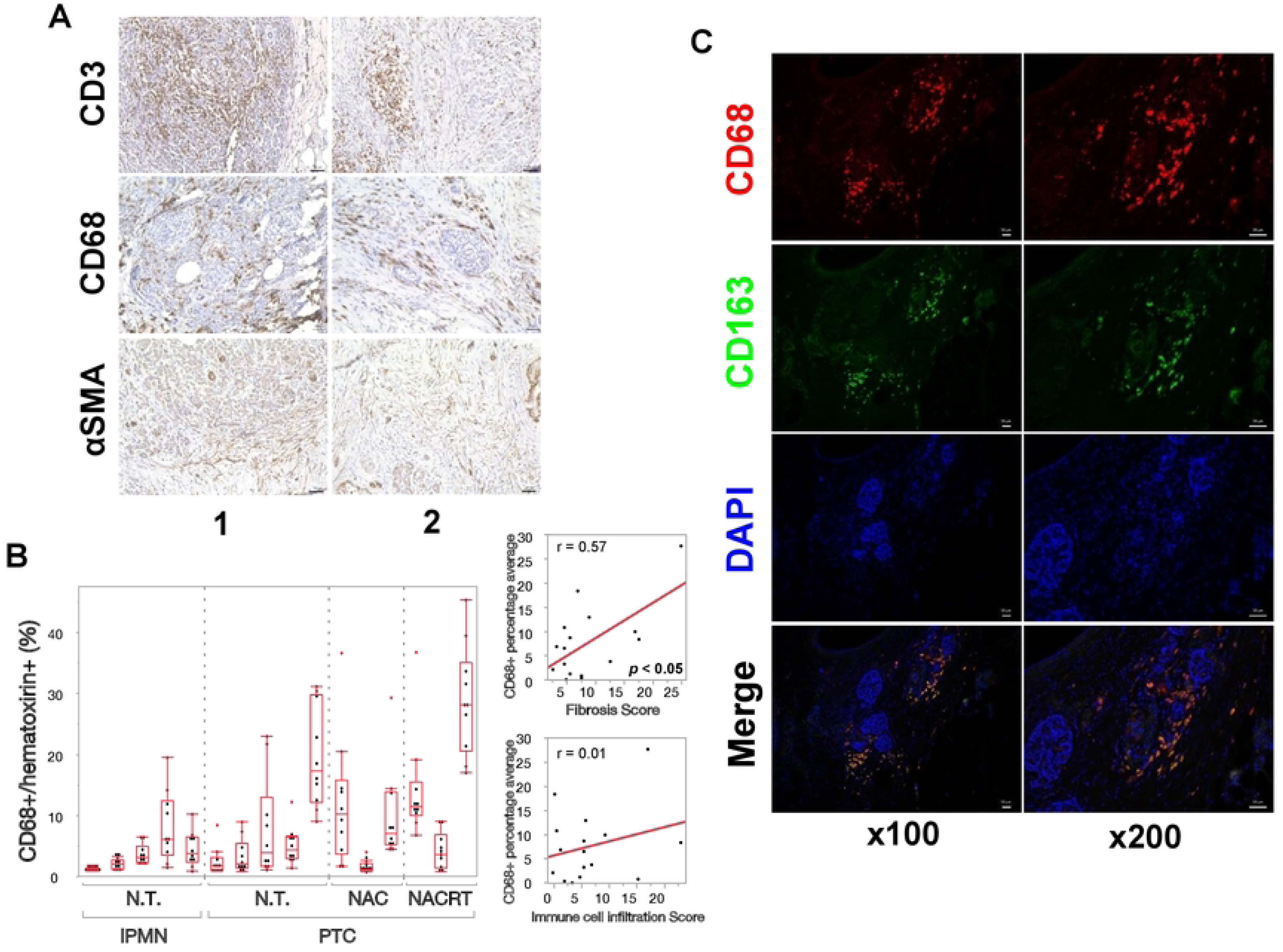
Macrophage-dominant inflammatory infiltrates in latent chronic pancreatitis A. High-magnification view of representative staining for major inflammatory cell markers. Scale bars = 50 µm B. Left: Mean CD68-positive scores for each case stratified by disease group. Bars indicate mean values. Right: Pearson correlations between CD68-positive scores ad either fibrosis scores or immune cell infiltration scores in the blinded evaluation. C. Double Immunofluorescence staining images of CD68 (red), CD163 (green), and DAPI (blue). Areas enclosed by yellow dotted boxes in the left images are shown at higher magnification on the right. Scale bars = 50 µm

Quantification of macrophages across all cases showed a positive correlation with the fibrosis score, but not with the overall immune cell infiltration score (total immune cell count). This suggests that macrophage levels may be regulated by mechanisms distinct from general inflammatory cell infiltration (Fig 2B). Furthermore, double immunofluorescence demonstrated abundant CD163-positive M2-type macrophages, even though the analyzed regions were located away from the primary tumor (Fig 2C).

### The role of *Enterococcus faecalis* in latent chronic pancreatitis

To clarify the involvement of *E. faecalis* in 16 cases of latent chronic pancreatitis, immunohistochemical analysis using polyclonal antibodies against Enterococcus species was performed (Fig 3A). No positive staining was observed in the pancreatic parenchyma any case; however, weak positivity was detected in the pancreatic ducts in 4 of 16 cases. Immunohistochemical staining for lipoteichoic acid (LTA), a cell wall component of gram-positive bacteria including *E. faecalis*, was negative in all cases. Genomic DNA was extracted from paraffin-embedded pancreatic tissue, and the gene encoding bacterial 16S rRNA was amplified. This analysis revealed the presence of bacterial nucleic acids in all cases, whereas no bands were detected by *E. faecalis*-specific PCR (Supplemental Fig 2A). In contrast, when genomic DNA was directly extracted from preserved frozen pancreatic tissue and the same analysis was repeated, *E. faecalis*-specific PCR was positive in some samples, however the positive rate differed depending on the sampling site even within the same case (S2B Fig).

**Fig 3.**
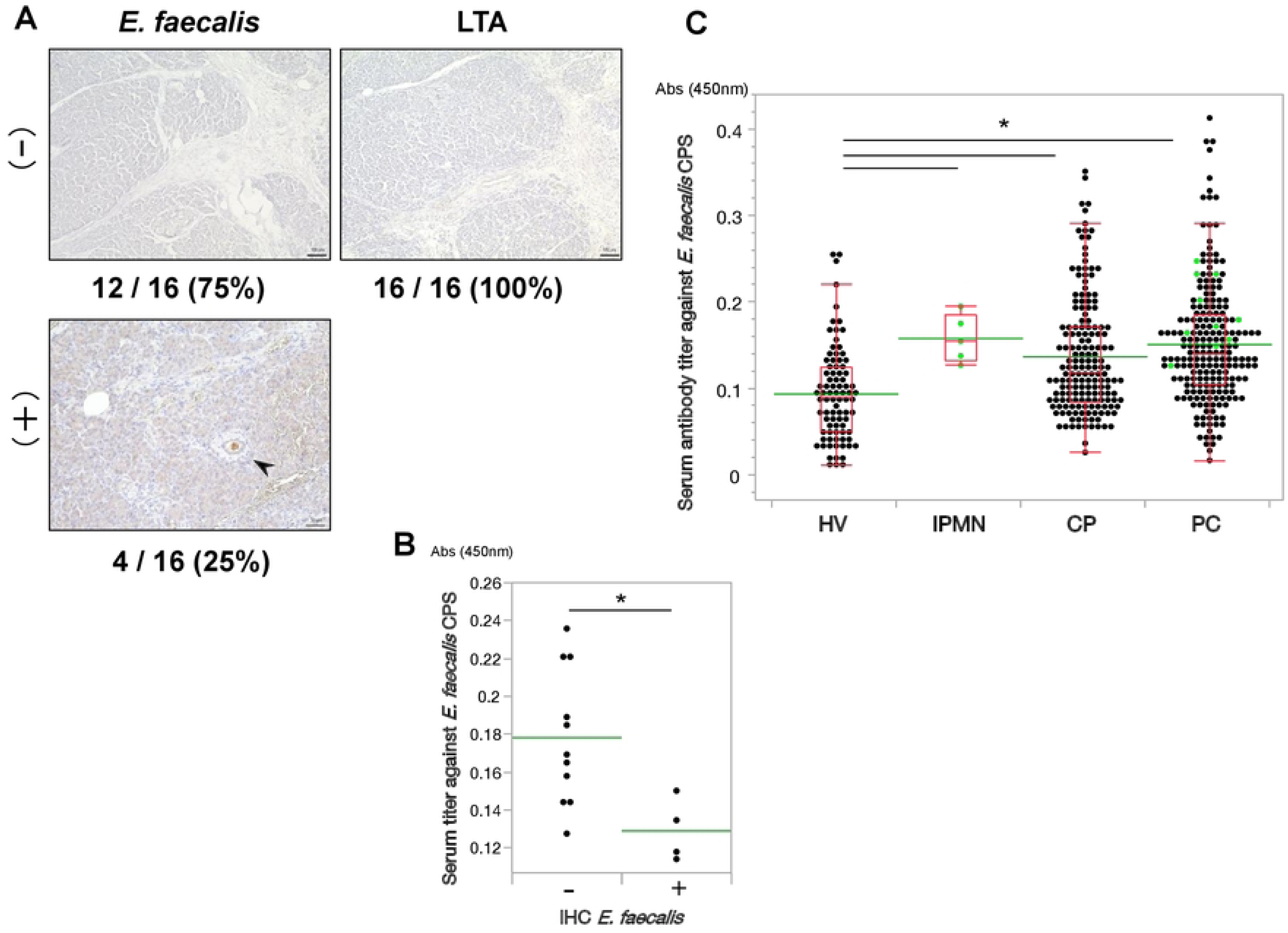
Role of Enterococcus faecalis in latent chronic pancreatitis A. Immunohistochemical staining of pancreatic tissue using anti-*E. faecalis* and anti-LTA antibodies. Scale bars = 50 µm B. Comparison of serum anti-E. faecalis CPS antibody titers according to *E. faecalis* immunostaining status in pancreatic tissue. Bars indicate mean values. *p < 0.05. C. Serum anti-*E. faecalis* CPS antibody titers in HV (healthy volunteers), IPMN (intraductal papillary mucinous neoplasm), CP (chronic pancreatitis), and PC (pancreatic cancer) evaluated by the Steel-Dwass test (*p < 0.05). Yellow-green plots indicate serum results from 15 of the 16 cases used for pancreatic histopathological analysis.

Next, we measured serum CPS antibody titers in these 16 cases. Cases showing weak *E. faecalis* positivity in the pancreatic duct exhibited significantly lower CPS antibody titers than immunohistochemically negative cases (Fig 3B). When CPS antibody titers in these 16 patients were compared with titers measured in stored sera from healthy volunteers, patients with chronic pancreatitis, and patients with pancreatic cancer, all 16 cases, including the IPMN cases, showed values exceeding the mean levels observed in chronic pancreatitis and pancreatic cancer (Fig 3C).

## Discussion

In our previous study, we examined autopsy pancreatic tissue and demonstrated that latent chronic pancreatitis frequently existed at sites corresponding to the primary location of pancreatic cancer development, rather than reflecting postmortem degeneration or obstructive pancreatitis secondary to cancer ^5^. In the present study, the positivity rate for Enterococcus species in pancreatic tissue was lower (25%) than that reported previously (50%). While positive staining for Enterococcus species was observed within the pancreatic duct, no cases exhibited strong, diffuse staining throughout the pancreas, as previously reported in two cases ^5^. To determine whether viable *E. faecalis* is present in the pancreas in vivo, we cultured macroscopically normal pancreatic tissue surrounding resected pancreatic cancer in six patients. Despite the fact that all patients underwent surgery and received perioperative antimicrobial agents, *Enterococcus casseliflavus*—a species of the genus Enterococcus—was isolated in one case (data not shown). Although these data are limited, the isolation of Enterococcus from pancreatic tissue despite antibiotic administration supports the notion that this bacterium is capable of colonizing the pancreas. In this study, serum CPS antibody levels were low in cases where Enterococcus species were histologically detected within the pancreatic ducts (Fig 3B). This suggests that the mere presence of bacteria in the pancreatic duct does not necessarily elicit an immune response ^16,17^. Rather, *E. faecalis* may induce pancreatitis accompanied by an immune response only when it invades the pancreatic parenchyma. As inflammation progresses and the pH of pancreatic juice decreases, *E. faecalis* may then be outcompeted or replaced by other bacteria. This hypothesis is supported by our previous report that *E. faecalis* is frequently detected when pancreatic function is preserved and pancreatic juice pH is maintained, but becomes undetectable when pancreatitis develops and the pancreatic juice pH can no longer be maintained at an alkaline level. ^18^. Integrating these observations with our previous findings—including those obtained from mouse experiments—we propose Fig 4 as the hypothetical model that best explains the data.

**Fig 4.**
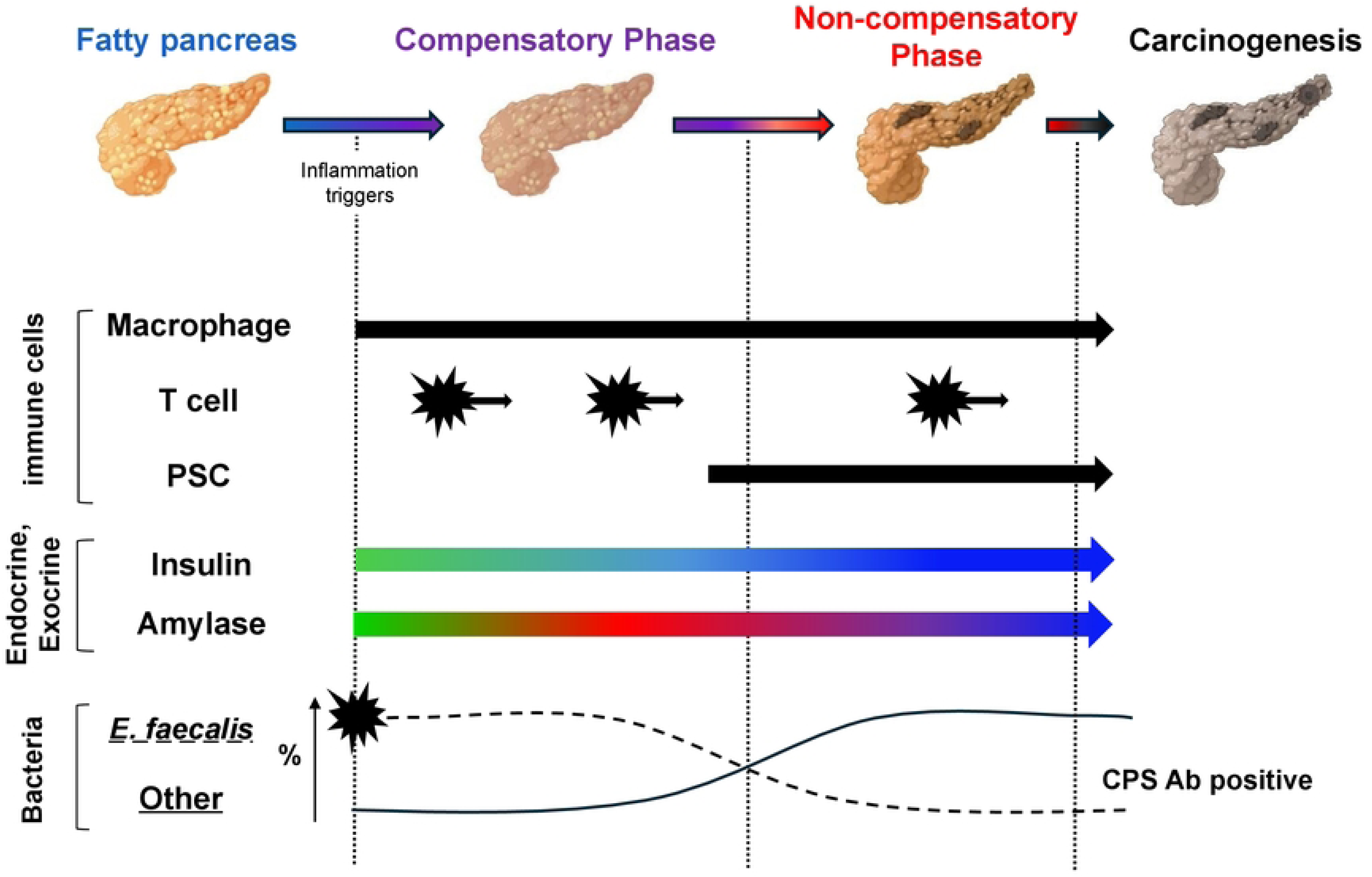
Hypothetical model of microbiota-driven pancreatic inflammation Hypothetical diagram illustrating functional alterations and dynamic changes in bacterial populations and immune cell subsets associated with the development of pancreatic inflammation.

The frequent and marked macrophage infiltration observed in pancreatic tissue from patients with latent chronic pancreatitis is noteworthy. One possibility is that macrophage numbers increase in response to stimulation by bacteria other than *E. faecalis* or by bacterial cellular components. Our findings differed from our initial expectations. Specifically, in cases where Enterococcus species were histologically detected within the pancreatic ducts, serum CPS antibody titers were low, and LTA immunostaining was negative in all instances. However, unlike in mouse experiments, it is impossible to track the temporal progression of *E. faecalis* infection in human cases. Therefore, we cannot rule out the possibility that it was difficult to detect traces of infection in the clinical samples obtained. While obtaining clinical samples suitable for analysis presents a significant challenge, we believe that analyzing a larger number of cases in the future will enable us to elucidate the underlying cause. Numerous studies have reported the presence of diverse bacterial populations in pancreatic cancer ^7,9,19,20^, indicating bacterial DNA or other components are present at least at the nucleic acid level. Analogous to *Helicobacter pylori* in gastric cancer ^21,22^, the failure to identify a single causative bacterium for pancreatic cancer within the pancreas may reflect a scenario in which long-term stimulation by *E. faecalis* induce persistent pancreatitis, which in turn drives a systemic antibody response. T cell infiltration in our study appeared highly localized. In chronic hepatitis, inflammatory responses occur frequently in specific lobular or portal regions ^23^, whereas in the pancreas, focal T cell accumulation may be reflecting site-specific immune responses occurring precisely where the cells cluster, for example around emerging atypical or pre-neoplastic cells.

While chronic gastritis can be diagnosed via endoscopy, chronic pancreatitis cannot. Although current serum biomarkers and imaging tests can narrow down the possibilities, it is believed that only a very small number of these cases will develop into pancreatic cancer. In the future, by combining analysis based on next-generation sequencing, screening using multi-biomarkers (including *E. faecalis* CPS), and tumor suppressor gene testing, it should be possible to identify individuals at high-risk for pancreatic cancer and thereby optimize strategies for surveillance and early intervention.

## Acknowledgement

We would like to thank to Dr Takashi Kumada at Gifu Kyoritsu University, Dr Toshifumi Ito at Osaka JCHO Hospital, and Dr Makoto Yamada at aMs new Ohtani Clinic for providing serum samples of pancreatic disease patients and healthy volunteers. The authors used a generative artificial intelligence tool (Perplexity, powered by GPT-5.1) to assist with English language editing and improvements in clarity of the manuscript. The tool was not used for data generation or analysis, and all scientific content was critically reviewed and approved by the authors.

---

**S1 Fig Immune cell distribution in pancreatic fibrosis**

A. Immunohistochemical staining images of major immune cell populations in pancreatic tissue. The left panels show regions at the parenchyma-fibrosis interface (1), and the right panels show densely fibrotic areas (2). Scale bar = 100 µm

B. Quantification of the proportion of antibody-positive cells among total cells in each region.

**S2 Fig Detection of bacterial genes in pancreatic tissue**

A. Agarose gel electrophoresis of PCR products targeting bacterial genes using DNA extracted from paraffin-embedded pancreatic tissue blocks used for histopathological analysis.

B. Agarose gel electrophoresis of PCR products targeting bacterial genes using DNA

extracted from frozen pancreatic tissue of pancreatic cancer patients. NT, non-tumor tissue; T, tumor tissue; PC, positive control, NC; negative control.

**S1 Table**

Clinical and pathological information for the 16 pancreatic tissue samples used for pathological analysis.

**S2 Table**

Primary and secondary antibodies used in this study

## Authors’ contribution

Kaho Nishikori: Investigation, Methodology, Formal analysis, Writing – original draft, review & editing, and Visualization.

Shinji Takamatsu: Methodology, Formal analysis, Writing – original draft, review & editing.

Munefumi Shimosaka: Investigation, Formal analysis, Writing – review & editing.

Risa Uemura: Investigation

Yudai Ishida: Investigation

Rikako Sugawa: Investigation

Muya Matsumoto: Investigation

Asuka Ogata: Investigation

Daisuke Sakon: Investigation, Formal analysis, Writing – review & editing.

Motoharu Inui: Resources

Daisaku Yamada: Resources

Hirofumi Akita: Resources

Jumpei Kondo: Formal analysis, Writing – review & editing.

Takahiro Kodama: Writing – review & editing.

Yoshihiro Kamada: Writing – review & editing.

Hidetoshi Eguchi: Resources, Writing – review & editing.

Eiichi Morii: Investigation, Writing – review & editing

Eiji Miyoshi: Conceptualization, Writing – original draft, review & editing, Supervision, Project administration, and Funding acquisition. All authors reviewed the manuscript.

